## Supplementary figures and images for "Impaired fatty acid import or catabolism in macrophages restricts intracellular growth of *Mycobacterium tuberculosis*"

### Figure 1 Supplement Figure 1

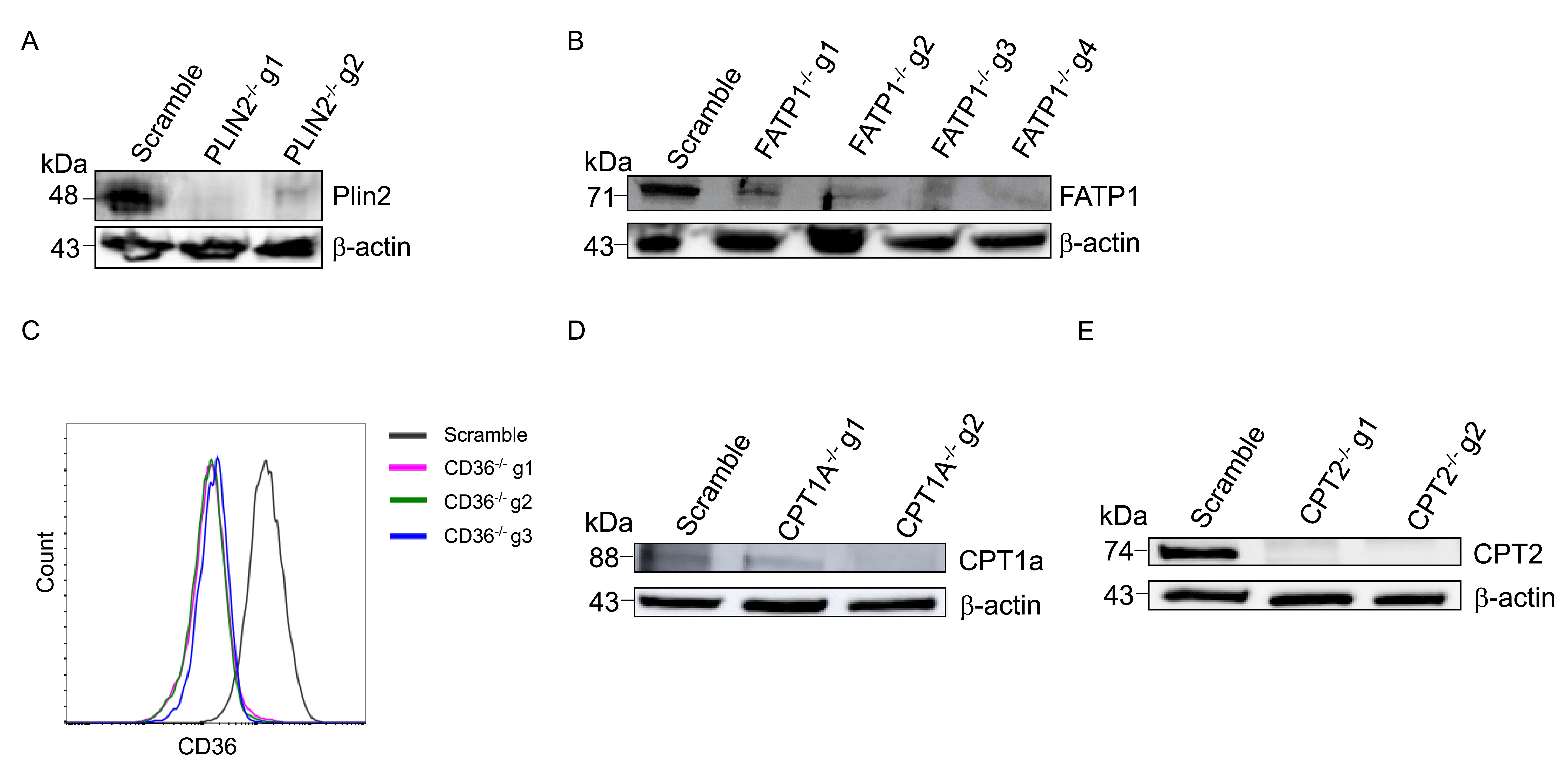

### Figure 2 Supplement Figure 1

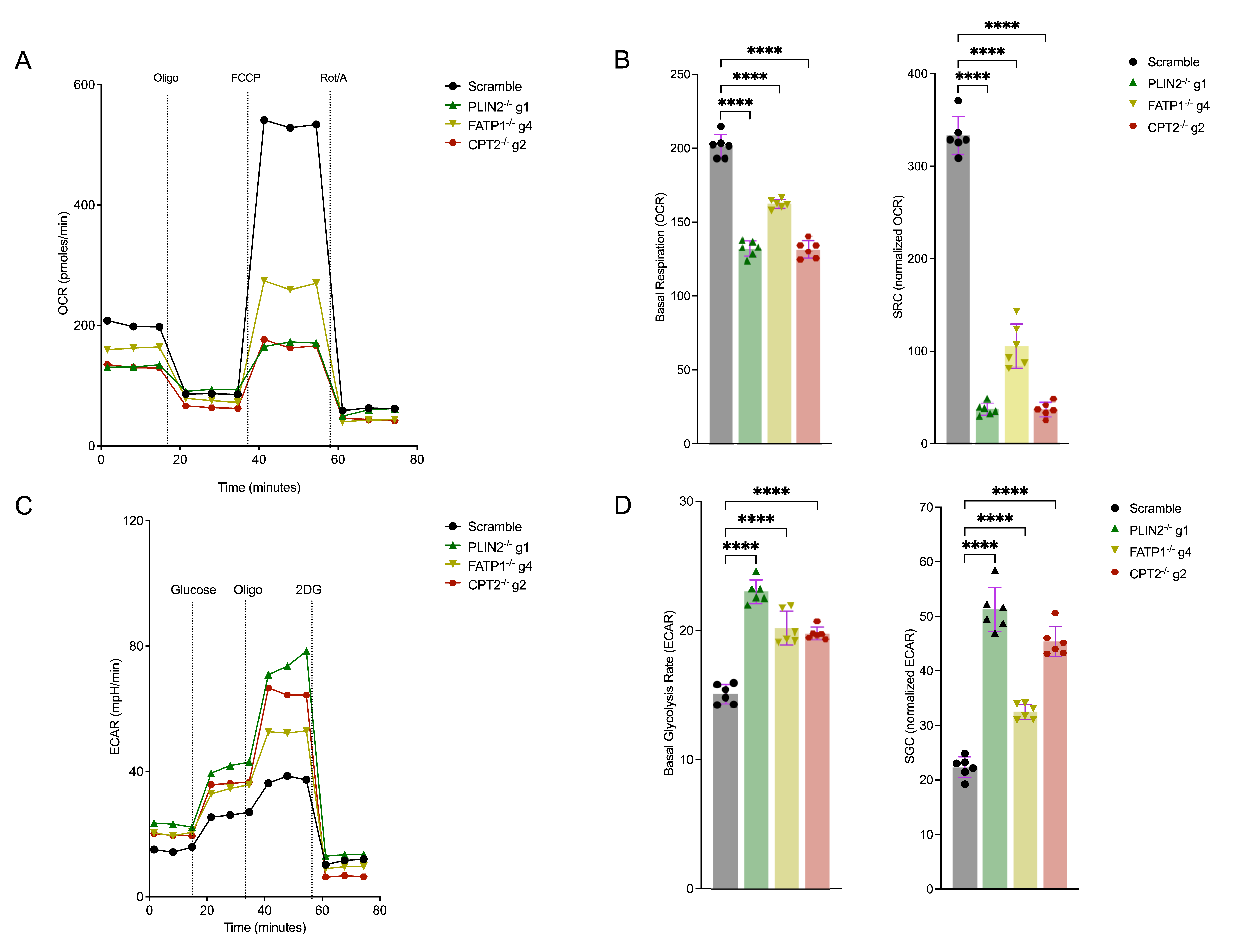

### Figure 2 Supplement Figure 2

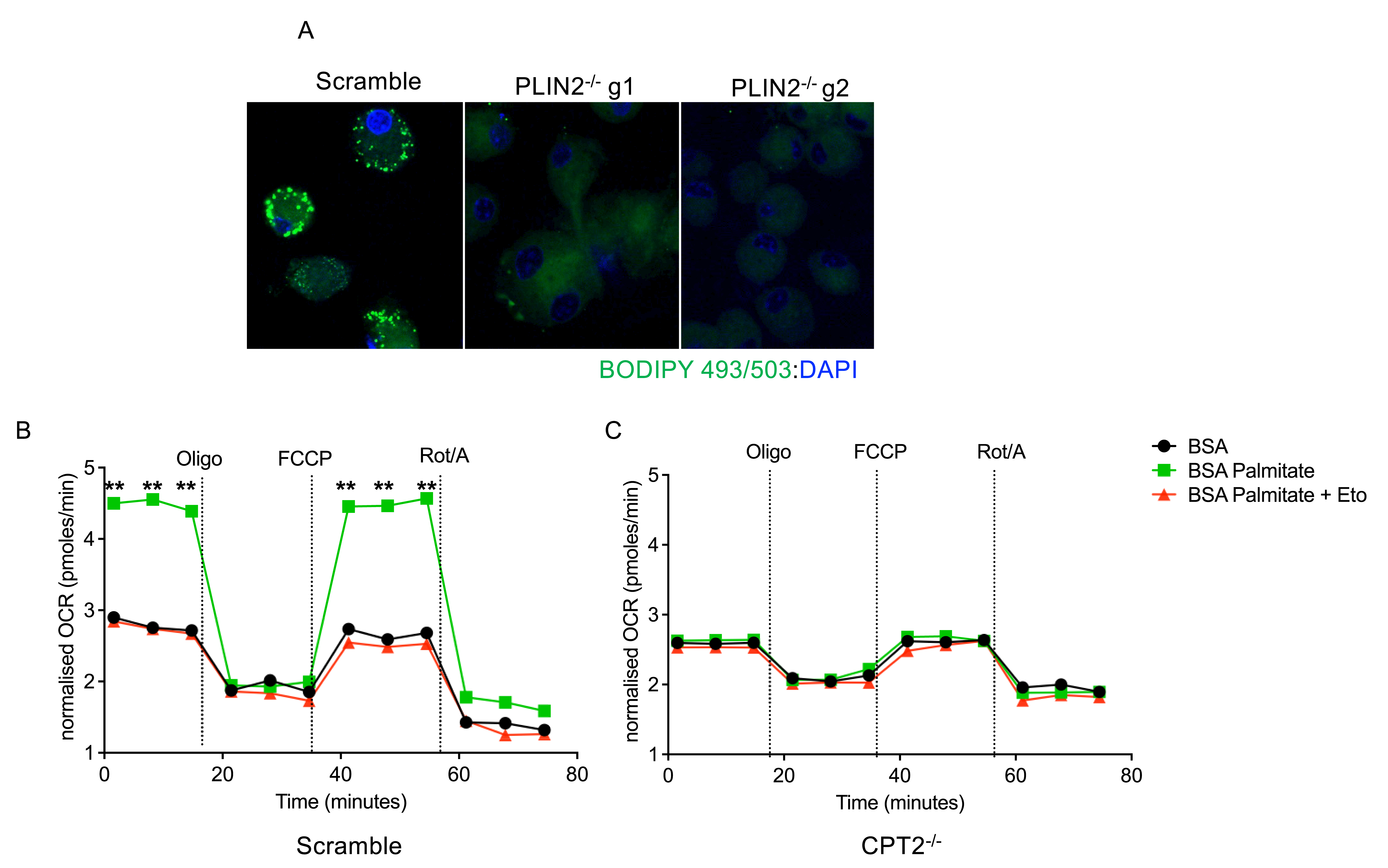

### Figure 2 Supplement Figure 2

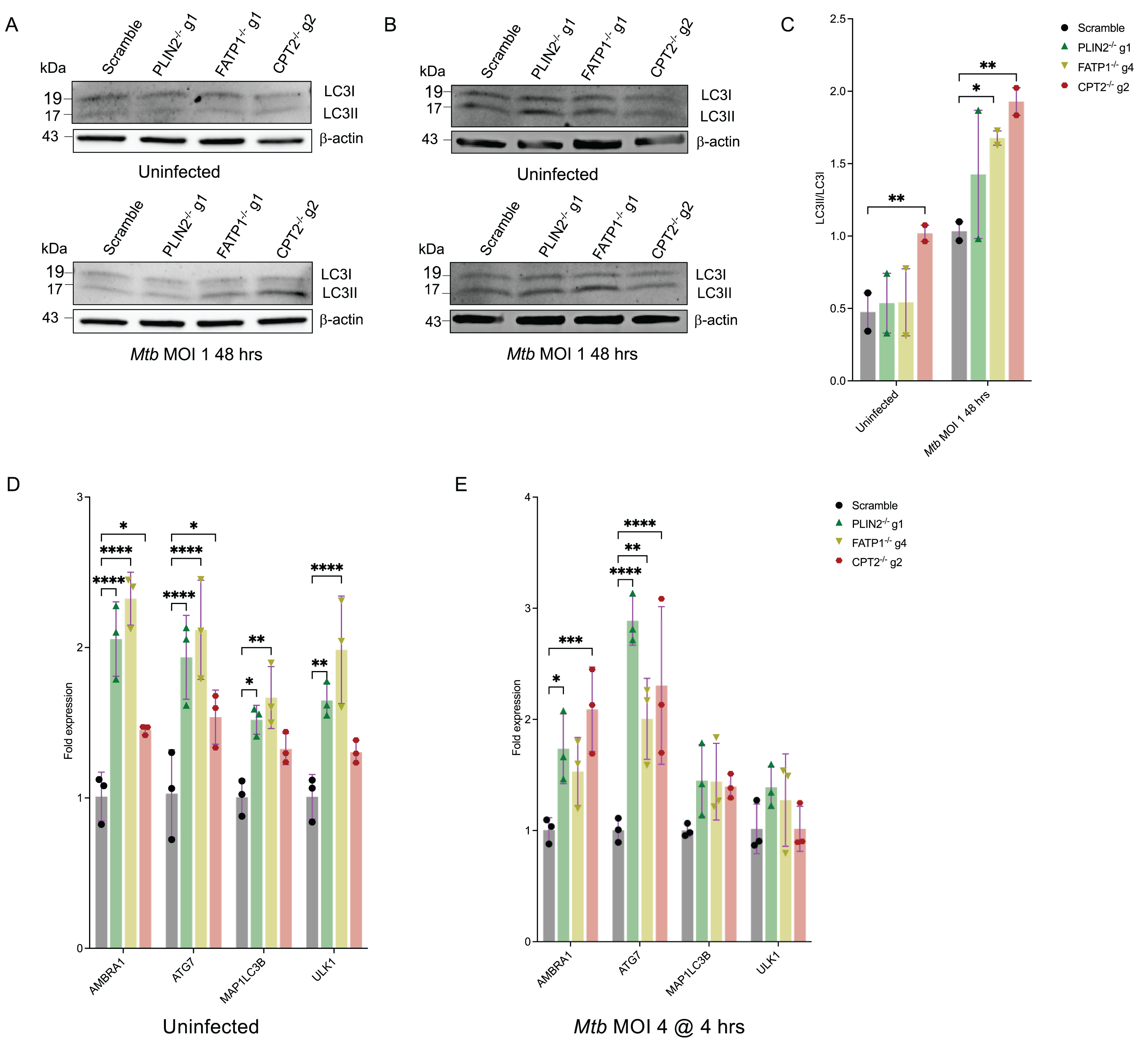

### Figure 3 Supplement Figure 1

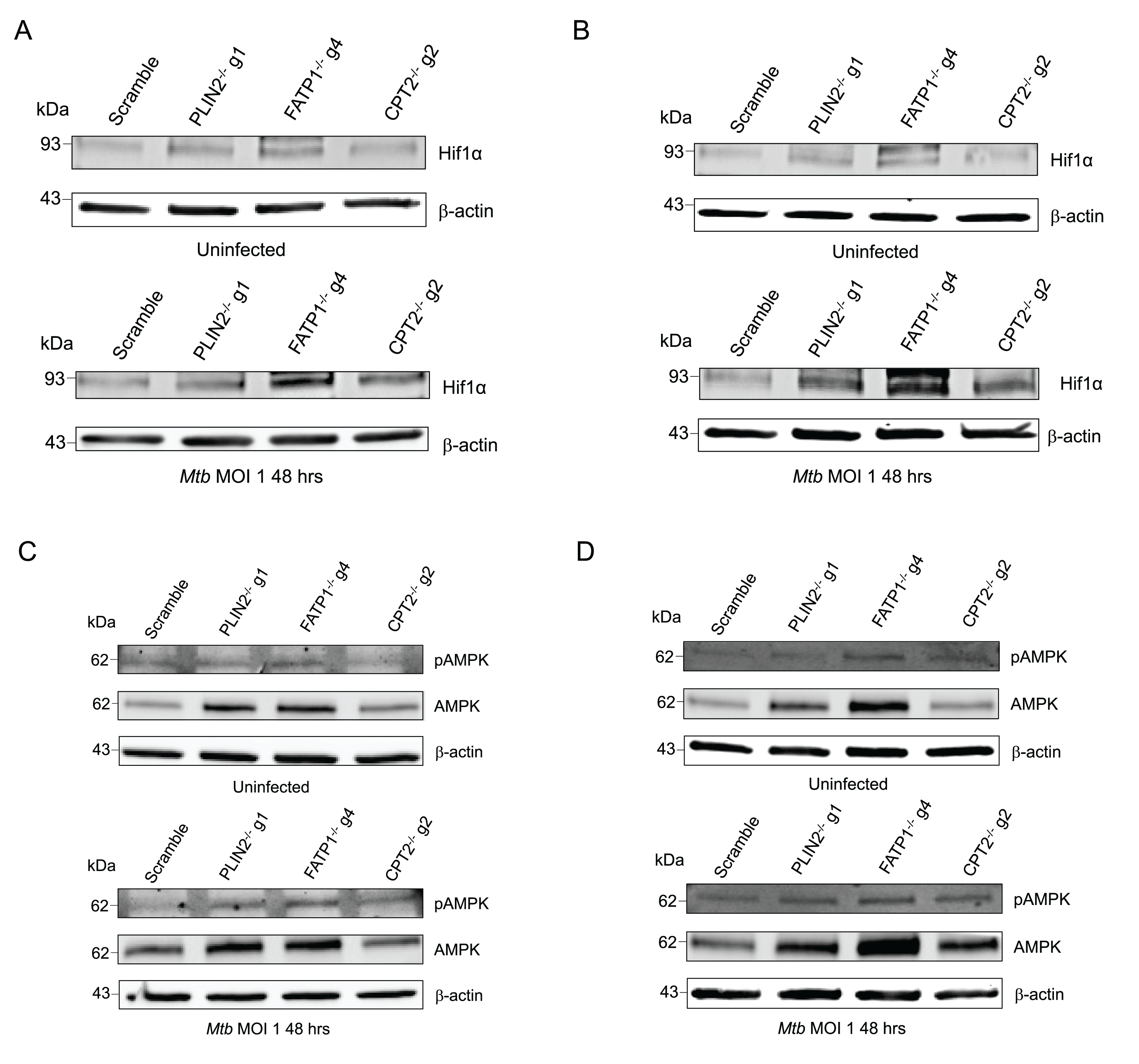

### Figure 6 Supplement Figure 1

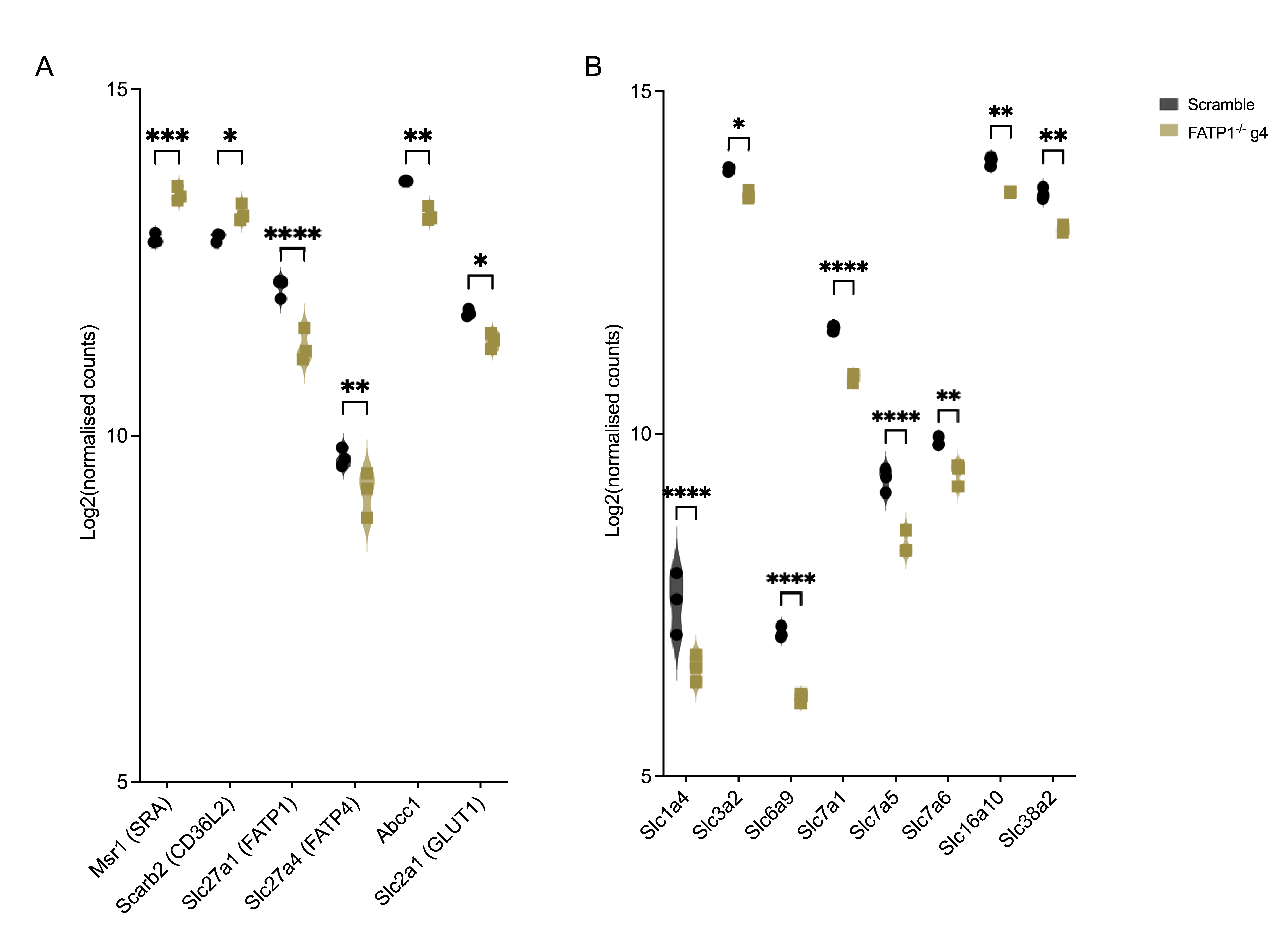

### Figure 6 Supplement Figure 2

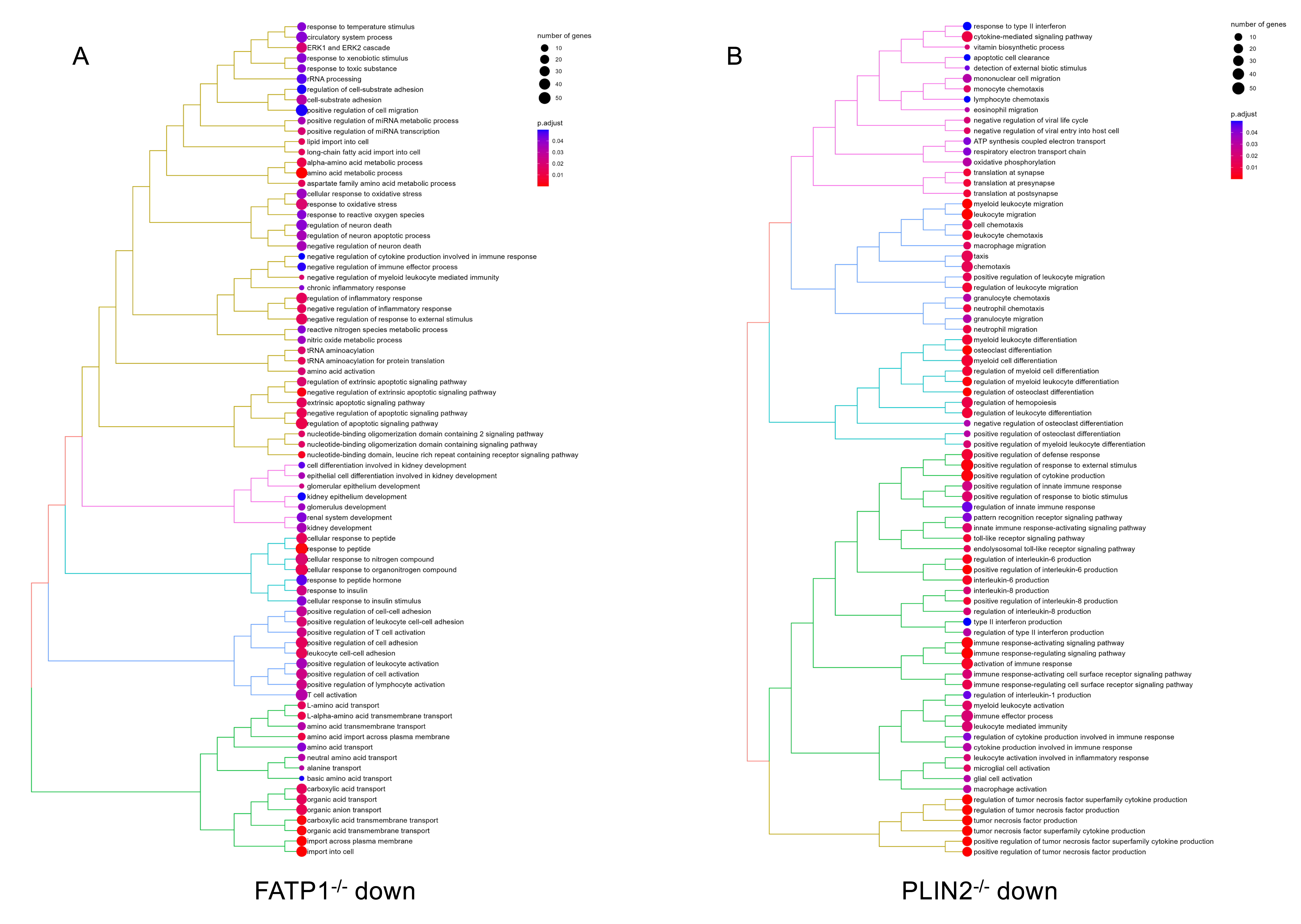

### Figure 6 Supplement Figure 3

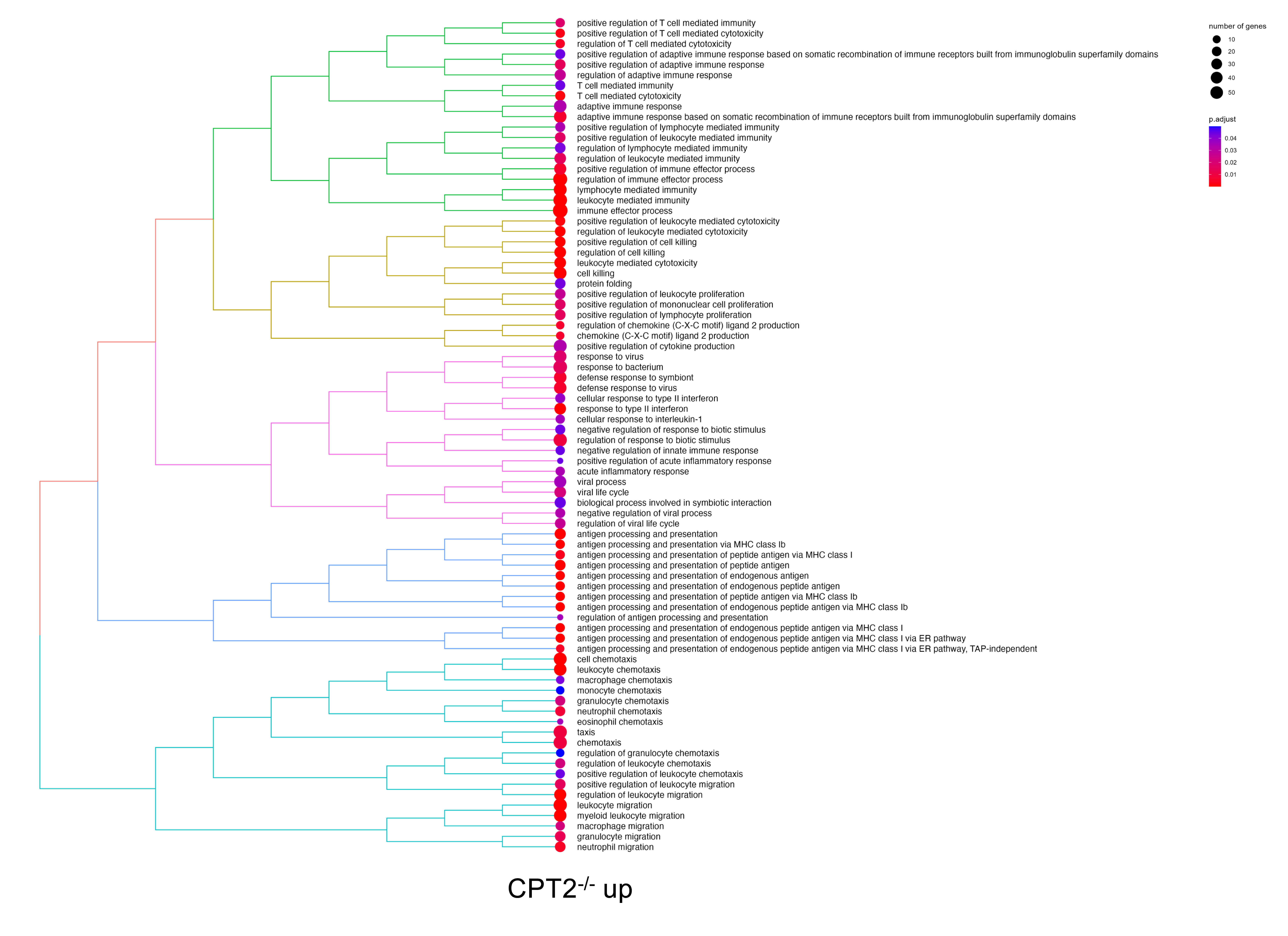

### Figure 6 Supplement Figure 4

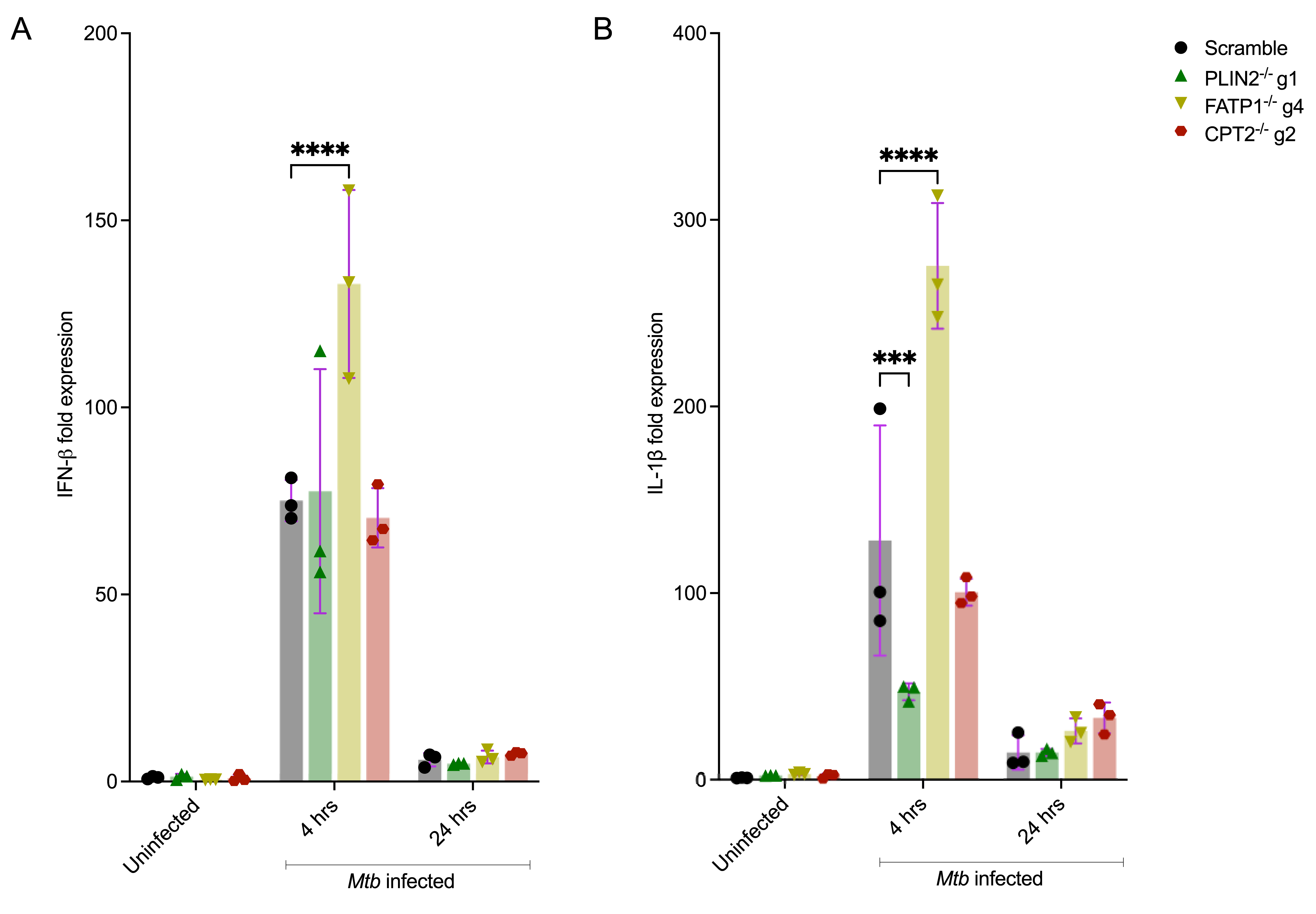

### Figure 7 Supplement Figure 1

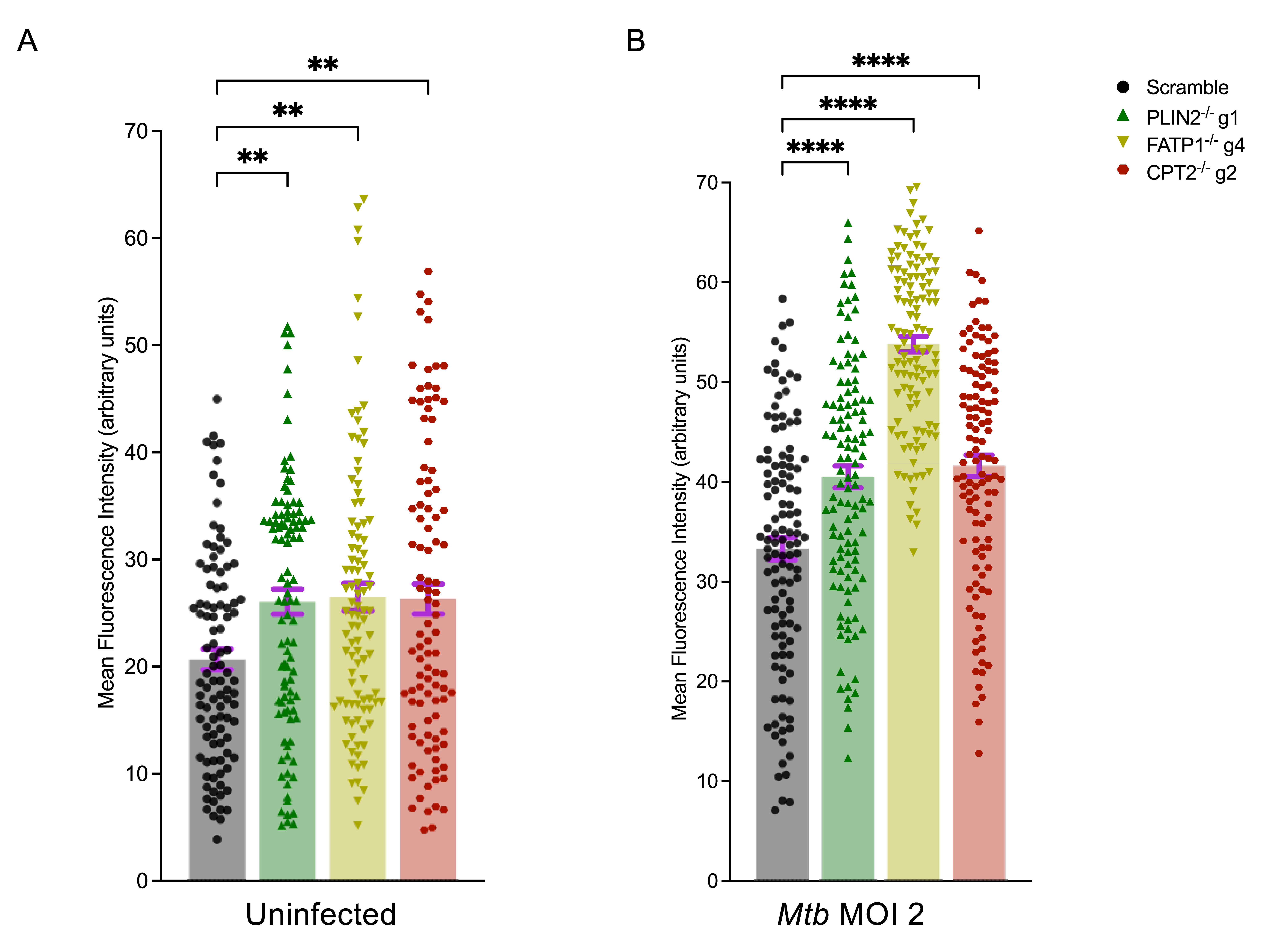
